# The Rab5-Rab11 endosomal pathway is required for BDNF-induced CREB transcriptional regulation in neurons

**DOI:** 10.1101/844720

**Authors:** Andrés G. González, Oscar M. Lazo, Francisca C. Bronfman

**Affiliations:** Department of Physiology, Faculty of Biological Sciences. Pontificia Universidad Católica de Chile, Santiago, Chile; Department of Neuromuscular Diseases, UCL Queen Square Institute of Neurology, University College London, London WC1N 3BG, UK; Center for Aging and Regeneration (CARE), Institute of Biomedical Sciences (ICB), Faculty of Medicine and Faculty of Life Sciences. Universidad Andrés Bello, Santiago, Chile

## Abstract

Brain-derived neurotrophic factor (BDNF) is a key regulator of the morphology and connectivity of central neurons. We have previously shown that BDNF/TrkB signaling regulates the activity and mobility of the GTPases Rab5 and Rab11, which in turn determine the post-endocytic sorting of signaling TrkB receptors. Moreover, altered Rab5 or Rab11 activity inhibits BDNF-induced dendritic branching. Whether Rab5 or Rab11 activity is important for local events only, or also for regulating nuclear signaling and gene expression, is unknown. Here, we investigated whether BDNF-induced signaling cascades were altered when early and recycling endosomes were disrupted by the expression of dominant negative mutants of Rab5 and Rab11. The activities of both Rab5 and Rab11 were required for sustained activity of Erk1/2 and nuclear CREB phosphorylation and for increased transcription of BDNF-dependent genes containing CRE-binding sites that include activity-regulated genes such as *Arc*, *Dusp1*, *c-fos* and *Egr1* and growth and survival genes such as *Atf3* and *Nf1*. Based on our results, we propose that the early and recycling endosomes provide a platform for the integration of neurotrophic signaling from the plasma membrane to the nucleus in neurons and that this mechanism likely regulates neuronal plasticity and neuronal survival.

**Significance Statement:** BDNF is a soluble neurotrophic factor that regulates plastic changes in the brain, including dendritic growth, by binding to its plasma membrane receptor TrkB. BDNF/TrkB activates signaling cascades leading to activation of CREB, a key transcription factor regulating circuit development and learning and memory. Our results uncover the cellular mechanisms that central neurons use to integrate the signaling of plasma membrane receptors with nuclear transcriptional responses. We found that the endosomal pathway is required for the signaling cascade initiated by BDNF and its receptors in the plasma membranes to modulate BDNF-dependent gene expression and neuronal dendritic growth mediated by the CREB transcription factor in the nucleus.

## Introduction

Neurotrophins shape the nervous system by regulating neuronal morphology and connectivity. Brain-derived neurotrophic factor (BDNF) is the most widely expressed member of this family in the brain and is associated with multiple functions, such as axon growth, dendritic branching, synaptic assembly and neurotransmission (Gonzalez et al., 2016; Kowianski et al., 2018). These diverse roles are explained by the multiplicity of signaling pathways triggered by BDNF. Upon binding of BDNF to tropomyosin-related kinase B (TrkB), the receptor is autophosphorylated at specific tyrosine residues in the intracellular domain, triggering the activation of the extracellular signal-regulated kinases 1 and 2 (ERK1/2) and the phosphatidylinositol 3-kinase (PI3K) pathways (Huang and Reichardt, 2003; Minichiello, 2009).

The signaling events triggered by BDNF/TrkB occur at the plasma membrane and regulate local events, such as protrusion of new neurites (Horch, 2004; Horch and Katz, 2002), or are propagated to the nucleus to impact gene expression (Cohen et al., 2011). Propagation and spatiotemporal dynamics of neurotrophin signaling largely depend on the trafficking of activated receptors as cargoes of membrane organelles known as ‘signaling endosomes’ (Barford et al., 2017; Villarroel-Campos et al., 2018). Consistently, endocytosis and trafficking of TrkB are required for activation of the PI3K signaling pathway and are necessary for sustained activation of Erk1/2 (Fu et al., 2011; Xu et al., 2016; Zheng et al., 2008). Interestingly, both mechanisms are required for BDNF-induced dendritic branching.

The intracellular trafficking of endocytic membranes is regulated by the Rab family of monomeric GTPases. Rab5 is a main determinant of early endosome identity; from there, cargoes can be sorted to Rab7-endosomes and lysosomes or recycled to the plasma membrane via Rab11-positive recycling endosomes (Goldenring, 2015; Jovic et al., 2010; Stenmark, 2009). We showed that TrkB localizes to both Rab5 and Rab11 endosomes after activation by BDNF in dendrites and cell bodies, and both GTPases are critical for BDNF-mediated dendritic branching (Lazo et al., 2013; Moya-Alvarado et al., 2018). While Rab11 has been shown to mediate BDNF/TrkB signaling events in dendrites and synapses (Huang et al., 2013; Lazo et al., 2013), to what extent they contribute to the propagation of BDNF-induced signaling cascades to the nucleus is unclear.

cAMP-response element-binding protein (CREB) is a transcription factor (TF) constitutively expressed by neurons and is known to mediate activity-dependent transcription, a process required for the development of neuronal connectivity. CREB exerts its effects by regulating the transcription of immediate-early genes, such as *c-fos*, *Arc*, *Bdnf* and the *Egr* family of TFs, among others. A signaling mechanism upstream of CREB includes the activation of neurotransmitter receptors and voltage gated channels that increase intracellular calcium levels (Flavell and Greenberg, 2008; Yap and Greenberg, 2018). Interestingly, CREB is also activated by BDNF signaling through TrkB receptors (Finkbeiner et al., 1997). Thus, CREB increases BDNF expression and, in turn, BDNF activates the phosphorylation of CREB, suggesting that there is a bidirectional positive feedback loop for CREB activation.

Different lines of evidence suggest that CREB activation by membrane receptors requires signaling from intracellular organelles. We previously demonstrated that trafficking of TrkB through the recycling pathway sensitizes hippocampal neurons to BDNF, increasing activation of the TF CREB at low ligand concentrations (Lazo et al., 2013). Furthermore, previous studies have indicated that PKA signaling from intracellular organelles, mediated by TSH receptors, is required to activate CREB in primary thyroid cells (Godbole et al., 2017). Finally, axonal signaling endosomes are required for CREB activation to induce survival in sympathetic neurons (Riccio et al., 1999; Yamashita and Kuruvilla, 2016)

Here, we hypothesized that disturbing the endosomal pathway by reducing the activity of Rab5 and Rab11 alters the temporal dynamics of BDNF-induced signaling pathways that regulate CREB activation and gene expression. Our results showed that the integrity of the early and recycling endosomes is crucial for CREB activation and further BDNF-induced transcriptional regulation and provided new evidence demonstrating that the endocytic route regulates plasma membrane to nuclear signaling in neurons.

## Results

### Sustained Erk1/2 activation requires Rab5 and Rab11 activity

Sustained signaling induced by neurotrophins is required for Erk1/2 nuclear translocation and differentiation of PC12 cells into a neuronal phenotype. However, transient activation of Erk1/2 retains it in the cytoplasm and promotes proliferation (Marshall, 1995). Consistently, sustained activation of Erk1/2 is crucial for structural plasticity in hippocampal neurons (Wu et al., 2001b). We used dominant negative (DN) mutants of Rab5 and Rab11 to investigate whether early and recycling endosomes play a role in sustained BDNF signaling. Hippocampal neurons (7 days *in vitro*, 7 DIV) were transduced with control EGFP, Rab5DN or Rab11DN adenovirus and were stimulated with BDNF 48 hours later for 5 minutes to evaluate activation of the signaling pathways or for 120 minutes to monitor sustained activation.

Five minutes of stimulation with BDNF led to robust phosphorylation of TrkB, Akt and Erk1/2, which was unaffected by the presence of the mutants Rab5DN and Rab11DN (Fig. 1A-B). Additionally, the levels of total TrkB, Akt and Erk1/2 remained unaltered after two days of expression of the mutants (Fig. 1A). However, after 120 minutes, the levels of phosphorylated Erk1/2 were still significantly increased in EGFP-expressing neurons, while cells expressing Rab5DN and Rab11DN showed levels that decreased to the basal levels (Fig. 1B). Akt activity showed a normal return to basal levels by 120 minutes of BDNF stimulation in control cells. Interestingly, neurons expressing the DN mutants showed even lower levels of Akt phosphorylation, suggesting that responsiveness to Akt can be progressively compromised when early and recycling endosomes are disrupted (Fig. 1B). No evident changes in the total levels of these kinases were found (Fig. 1A).

**Fig. 1.**
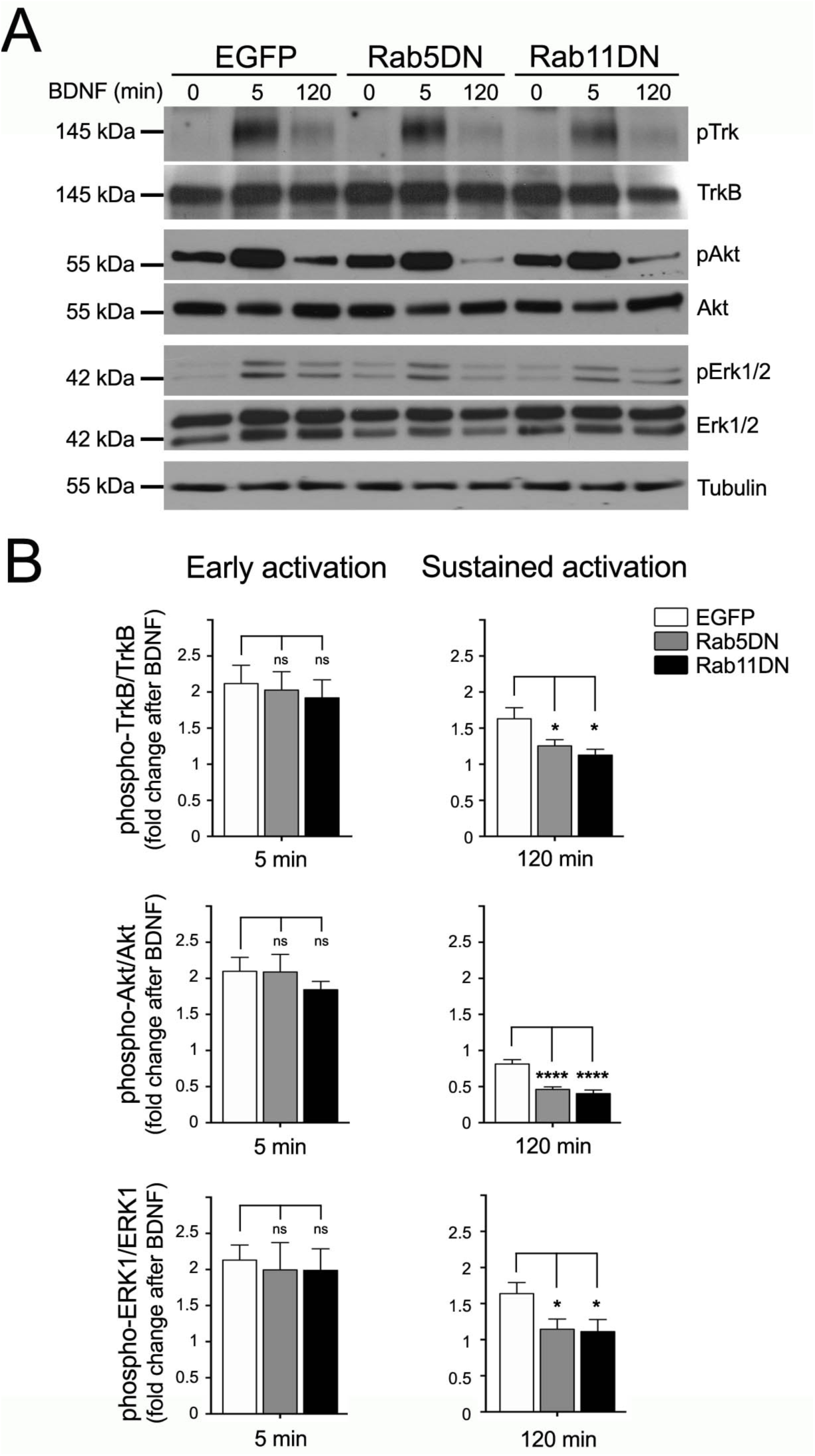
Rab5 and Rab11 activity is required for sustained BDNF-downstream signaling. Hippocampal neurons expressing control EGFP or dominant negative (DN) mutants of Rab5 and Rab11 were stimulated with BDNF for 0, 5 or 120 minutes, and lysates were probed for total and phosphorylated TrkB (Y515), Akt (S473) and Erk1/2 (T202/Y204). Representative western blots (**A**) and quantification of 5–7 independent experiments (**B**) are shown. Differences between 5 and 120 minutes after BDNF stimulation are indicated in brackets. One-way ANOVA followed by Brown-Forsythe and Welch post-test for multiple comparison. *, p < 0.04; ****, p < 0.0001; and NS, non-significant.

To investigate how sustained activation of Erk1/2 was affected at specific neuronal compartments, we performed immunofluorescence to detect phosphorylated Erk1/2 at T202 and Y204 (pErk1/2). The treatment of neurons with BDNF increased the pErk1/2-associated fluorescence in the nuclei, cell body and dendrites of hippocampal neurons (Fig. 2A). The staining was abolished in the somas and nuclei when cells were cotreated with BDNF and the Erk1/2 inhibitor PD98059 (Fig. 2A-C), demonstrating the specificity of the staining. Consistent with the results of Fig. 1, neurons expressing Rab5DN or Rab11DN and stimulated with BDNF for 60 minutes showed a decreased response to BDNF (Fig. 2D and E). Fluorescence intensity was quantified in dendrites, somas and nuclei, and expression of Rab5DN significantly reduced pErk1/2-associated fluorescence in all these neuronal compartments. Although the effect of Rab11DN in dendrites was unclear, the activity of Erk1/2 in the soma was significantly decreased and completely eliminated in the nucleus by this mutant (Fig. 2D and E). Together, these preferential effects on the nuclear signaling predict a stronger effect on gene regulation compared to local signaling from the plasma membrane. In summary, our results suggest that although activation of TrkB, Akt and Erk1/2 can occur normally in neurons expressing DN mutants, sustained signaling by Erk1/2 requires the activity of the small GTPases Rab5 and Rab11, potentially resulting in effects at the transcriptional level.

**Fig. 2.**
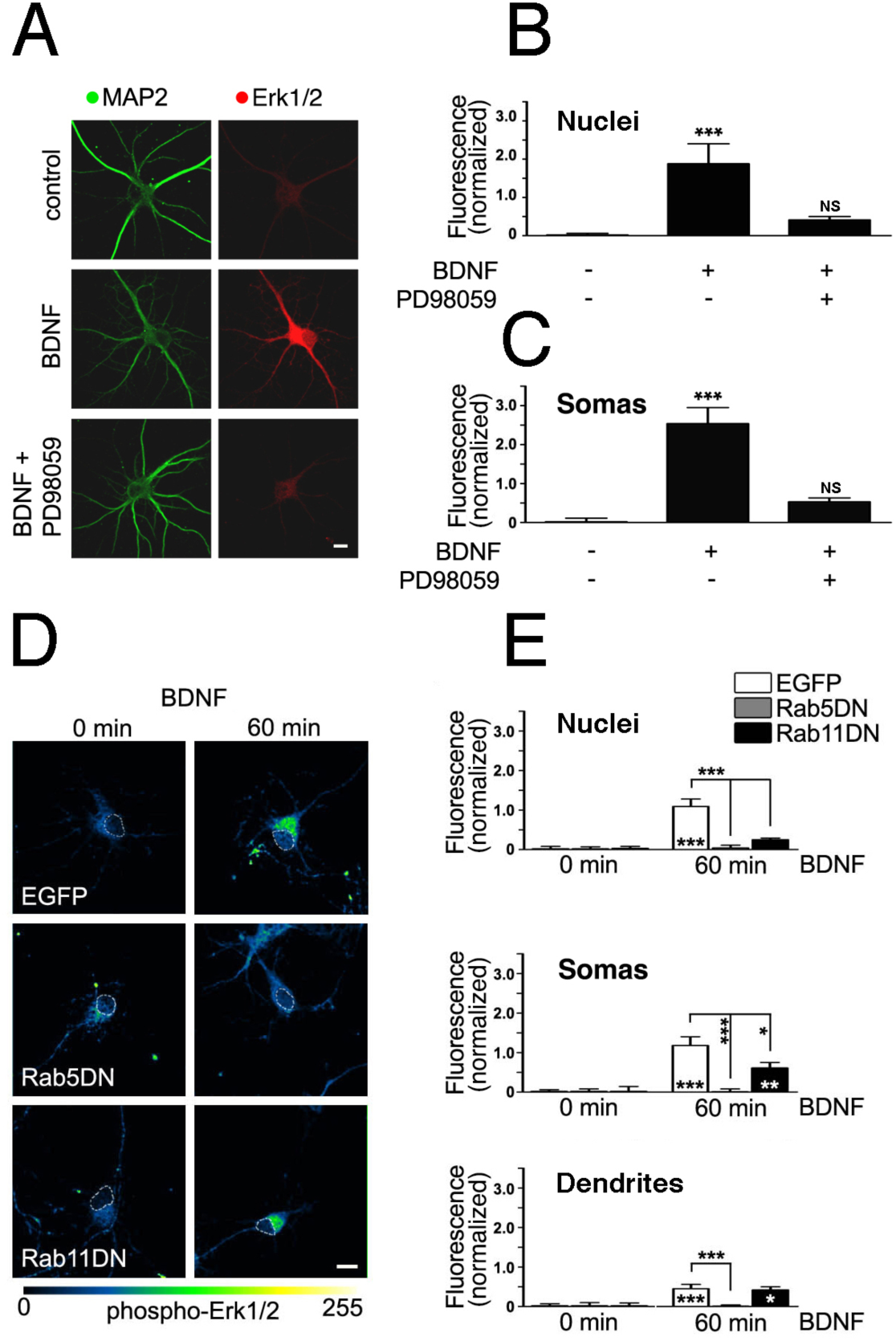
Rab5 and Rab11 activity is required for sustained BDNF-Erk1/2 signaling. The ability of PD98059 to inhibit BDNF-induced phosphorylation of Erk1/2 was confirmed by stimulating hippocampal neurons with BDNF for 15 minutes and performing double immunofluorescence for MAP2 and pErk1/2 (**A**). The ability of PD98059 to reduce pErk1/2 staining was evident in somas and nuclei, and the quantifications are shown in (**B**-**C)**. Scale bar: 10 µm. Hippocampal neurons expressing control EGFP or dominant negative (DN) mutants of Rab5 and Rab11 were stimulated with BDNF for 0 o 60 minutes, and sustained activation of Erk1/2 was analyzed in different neuronal compartments (nuclei, somas, dendrites) by immunofluorescence (**D**). Nuclei were traced using Hoechst staining as a reference and are indicated here with a dashed line. Scale bar: 10 µm. Quantification of the normalized intensity from 30 nuclei and somas and 60 dendrites from 3 independent experiments is shown (**E**). In D, differences between time=0 and 60 minutes of BDNF stimulation are indicated at the bottom of the bar. The effects of the expression of the Rab5 and Rab11 mutants compared with EGFP after 60 minutes of BDNF stimulation are indicated in brackets. Statistical differences were tested by using one-way ANOVA, followed by Dunnett’s post-test for multiple comparison. *, p < 0.05; **, p < 0.01; ****, p < 0.0005; and NS, non-significant.

### Rab5 and Rab11 activity is required for BDNF-induced CREB phosphorylation

To investigate whether activation of the transcription factor CREB downstream of BDNF signaling was also regulated by Rab5 and Rab11, we transduced 7 DIV hippocampal neurons with control EGFP, Rab5DN or Rab11DN adenoviral vectors. After 48 hours, cells were stimulated with BDNF for 15 minutes and compared to non-stimulated controls, and the levels of phosphorylated CREB at the S133 residue were evaluated by western blotting (Fig. 3A). EGFP-expressing neurons responded to BDNF stimulation with a significant increase in pCREB levels, while expression of DN mutants of Rab5 and Rab11 completely blocked this effect (Fig. 3A and B). To confirm this effect and further investigate the spatiotemporal dynamics of CREB activation, we used confocal microscopy to evaluate phosphorylated CREB in the nuclei of neurons by immunofluorescence. Neurons expressing control EGFP or the DN mutants of Rab5 and Rab11 were stimulated for 0, 15, 30 or 60 minutes with BDNF. While control EGFP expressing neurons showed a strong increase in CREB activation after 15–60 minutes of stimulation (Fig 3C-F), the expression of Rab5DN and Rab11DN significantly blocked BDNF-induced activation of CREB in the nucleus (Fig 3C-F), consistent with the effects on nuclear pErk1/2. These results suggest that BDNF/TrkB signaling requires Rab5 and Rab11 activity to induce nuclear CREB activation.

**Fig. 3.**
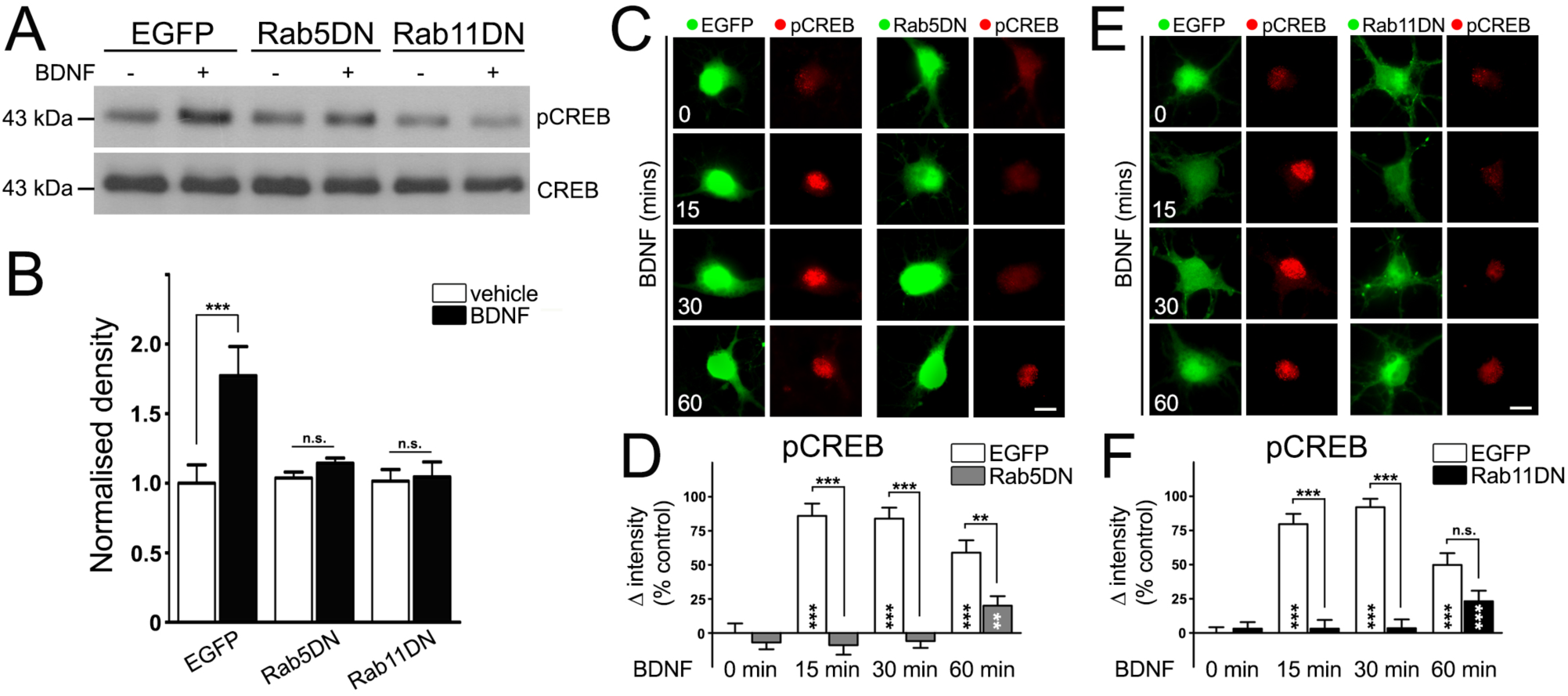
Nuclear CREB phosphorylation requires Rab5 and Rab11 activity. Hippocampal neurons expressing control EGFP or dominant negative (DN) mutants of Rab5 and Rab11 were stimulated with BDNF for 15 minutes, and lysates were probed for total and phosphorylated CREB. Representative western blots (**A**) and quantification of 4 independent experiments (**B**) are shown. Statistically significant differences between vehicle and BDNF treatments analyzed by using t-tests are indicated. To specifically study the amount of phosphorylated CREB in the nucleus, we performed stimulation with BDNF for 0, 15 or 60 minutes, and we detected phosphorylated CREB by immunofluorescence (**C-E**). Quantification of the normalized intensity from 42–112 nuclei from 4 independent experiments is shown (**D-F**). Datasets were tested by using one-way ANOVA, followed by Bonferroni’s post-test for multiple comparison. Statistically significant differences in pCREB at different time points compared to time = 0 minutes are indicated within the bar. Differences between EGFP controls and mutant-expressing neurons at specific time points are indicated in brackets. Significance levels are labelled as follows: **, p < 0.01; ***, p < 0.005; and NS, non-significant. Scale bars: 10 µm.

### CREB phosphorylation is regulated by the Erk1/2 pathway and is required for BDNF-induced dendritic branching

In different models, CREB can be activated through the Ras-Erk and PI3K-Akt signaling pathways (Du and Montminy, 1998; Finkbeiner et al., 1997). Since both signaling pathways are downstream of BDNF/TrkB, we decided to evaluate their contribution to the activation of CREB in our system. To achieve this aim, we used 9 DIV hippocampal neurons treated with a selective inhibitor of Erk1/2, PD98059, (Murray et al., 1998) or a potent general inhibitor of PI3Ks, LY294002 (Leemhuis et al., 2004), and we stimulated these cells with BDNF for 15 minutes. Activation of CREB in the nucleus was measured by using immunofluorescence (Fig. 4A-B). BDNF stimulation induces a strong increase in nuclear pCREB, which is blocked by the Trk inhibitor K252a (Tapley et al., 1992). Neurons treated with PD98059 showed robust inhibition of Erk1/2 phosphorylation in both the somas and nuclei (Fig. 2A-C), which, in turn, fully inhibited CREB activation in the nucleus (Fig. 4A-B). By contrast, the presence of LY294002 did not alter the BDNF-induced activation of CREB, suggesting that the PI3K-Akt cascade is dispensable to induce CREB activation in hippocampal neurons. To monitor whether LY294002 interfered with the activation of Akt without any effect on TrkB, we examined phosphorylation of both proteins by western blotting (Fig. 4C). We confirmed that although BDNF-dependent Trk phosphorylation was independent of the presence of the inhibitor, Akt phosphorylation at serine 473 was completely blocked. Therefore, BDNF-induced nuclear phosphorylation of CREB essentially depends on Erk1/2 activation in our model.

**Fig. 4.**
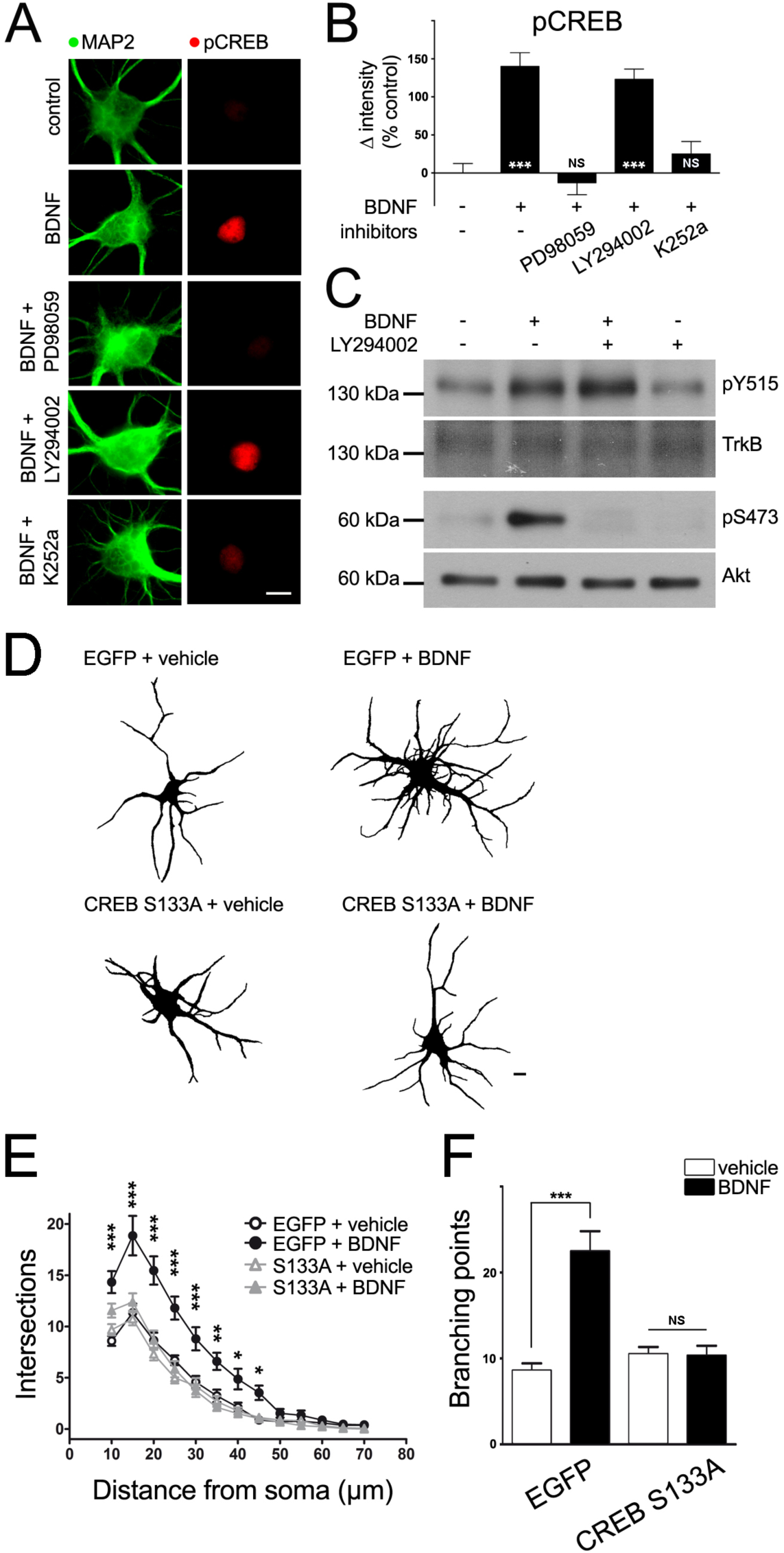
Phosphorylation of CREB is downstream of the Erk1/2 pathway and is required for BDNF-induced dendritic branching. Hippocampal neurons were stimulated with BDNF for 15 minutes in the presence of the MEK1/2 inhibitor PD98059, the PI3K inhibitor LY294002 or the Trk kinase activity inhibitor K252a and then probed for phosphorylated CREB and MAP2 by immunofluorescence (**A**). Quantification of 30 nuclei per condition from 3 independent experiments showed a complete block of BDNF-induced CREB phosphorylation in the presence of PD89059 (**B**). Significant differences compared to non-stimulated neurons were analyzed by using one-way ANOVA, followed by Bonferroni’s test, and are indicated within the bars. The dependence on TrkB was confirmed by the robust inhibition of BDNF-induced CREB phosphorylation by K252a. By contrast, LY294002 did not have any effect on pCREB, despite its significant inhibition of Akt phosphorylation (pS473), as shown in **C**. Null effect of LY294002 on TrkB phosphorylation (pY515) is also shown. Quantification of the pCREB levels confirmed the inhibition in the nuclei (**E**) and somas (**F**). Statistically significant differences were analyzed by using one-way ANOVA and Bonferroni’s post-test for multiple comparison and are indicated over the respective bars (n= 3 independent experiments; 20 neurons per condition). To confirm whether Erk1/2-mediated phosphorylation of CREB at S133 was critical for a cellular response, we compared BDNF-induced dendritic branching in neurons expressing either EGFP or a non-phosphorylatable mutant of CREB (S133A). Immunostaining for MAP2 was used to analyze the morphology of the dendritic arbor (**G**). Sholl’s analysis (**H**) and direct counting of the number of branching points (**I**) from 18 neurons per condition in 3 independent experiments showed that phosphorylation of CREB is required for BDNF-induced dendritic branching. Sholl’s curves were compared using two-way repeated-measures ANOVA, followed by Bonferroni’s post-test for multiple comparison. Branching point data were analyzed by using Student’s t test. Significance levels for the different statistical tests are labelled as follows: *, p < 0.05; **, p < 0.01; ***, p < 0.005; and NS, non-significant. Scale bars: 10 µm.

We previously established that Rab5 and Rab11 play a critical role in BDNF-induced dendritic branching (Lazo et al., 2013; Moya-Alvarado et al., 2018), and we showed that both GTPases are required for BDNF-induced CREB phosphorylation (Fig. 3). In addition, BDNF/TrkB signaling regulates Rab5 and Rab11 activity and dynamics, suggesting that both GTPases are downstream targets of BDNF/TrkB signaling (Lazo et al., 2013; Moya-Alvarado et al., 2018). To further corroborate that CREB is required for BDNF-induced dendritic branching, we expressed a non-phosphorylatable CREB mutant (CREB S133A) in 7 DIV hippocampal neurons and stimulated them with BDNF (Fig. 4D-F). Since the expression of CREB S133A for 48 hours did not decrease the complexity of dendritic arbors, we concluded that it does not affect basal dendritic morphology at this time point. However, the BDNF-induced increase in complexity, as measured by using Sholl’s analysis (Fig. 4E), and, more specifically, the increase in the number of branches (Fig. 3F) were completely abolished by the presence of the CREB S133A mutant, suggesting that CREB is needed for BDNF-dependent dendritic branching downstream of TrkB and Erk1/2. Taken together, our results suggest that BDNF/TrkB signaling activates nuclear CREB through the Erk1/2 pathway, requiring an increase in the activities of Rab5 and Rab11 to regulate gene expression.

### Rab11 is required to induce CREB-mediated gene expression after BDNF stimulation

We have previously shown that Rab5 activity is required for both the maintenance of basal dendritic complexity and for the increase in dendritic complexity induced by BDNF in hippocampal neurons. Rab11DN-expressing neurons, however, did not show a decrease in basal dendritic complexity at 7 DIV and are thus specifically required for BDNF-induced dendritic branching (Lazo et al., 2013; Moya-Alvarado et al., 2018). These results indicated that Rab5 has more pleiotropic effects on dendritic morphology than Rab11 does. During the endocytic sorting of signaling receptors, Rab11 recycling processes are downstream of Rab5-dependent early endosomal sorting. Normally, receptors are internalized to Rab5 early endosomes and then sorted to either Rab11 recycling endosomes or Rab7 late endosomes (Bronfman et al., 2014; Jovic et al., 2010). For these reasons, we decided to negatively regulate the activity of Rab11 (and not Rab5) to further study the effect of specifically altering the Rab11 recycling endosomal route on cAMP-response element (CRE)-dependent gene expression induced by BDNF since CREB exerts its function by binding specifically to genes targeted by CRE regulatory elements (Cha-Molstad et al., 2004).

Figure 3 shows that reducing Rab5 or Rab11 activity significantly decreased BDNF-induced CREB phosphorylation. However, according to previous reports, despite being a key step, serine-133 phosphorylation of CREB may be insufficient to trigger gene transcription (Finsterwald et al., 2010; Zhang et al., 2005). To further address whether the recycling endosomes regulated by concerted activities of Rab5 and Rab11 are required for CRE-dependent gene expression induced by BDNF, we first studied the effect of downregulating the activity of Rab11 on the levels of the activity-regulated cytoskeleton-associated protein (*Arc*) gene by qPCR. *Arc* is an immediate early gene and a well-known transcriptional target of BDNF that possesses CREs (Kawashima et al., 2009). We used 7 DIV hippocampal neurons expressing EGFP or Rab11DN for 2 days and then stimulated the neurons with BDNF for 4 hours. BDNF increased *Arc in* EGFP-expressing neurons by 5.4-fold (Fig. 5A), while Rab11DN-expressing neurons displayed an attenuated response (3.3-fold), suggesting that Rab11 significantly mediates transcriptional response to neurotrophic signaling.

**Fig. 5.**
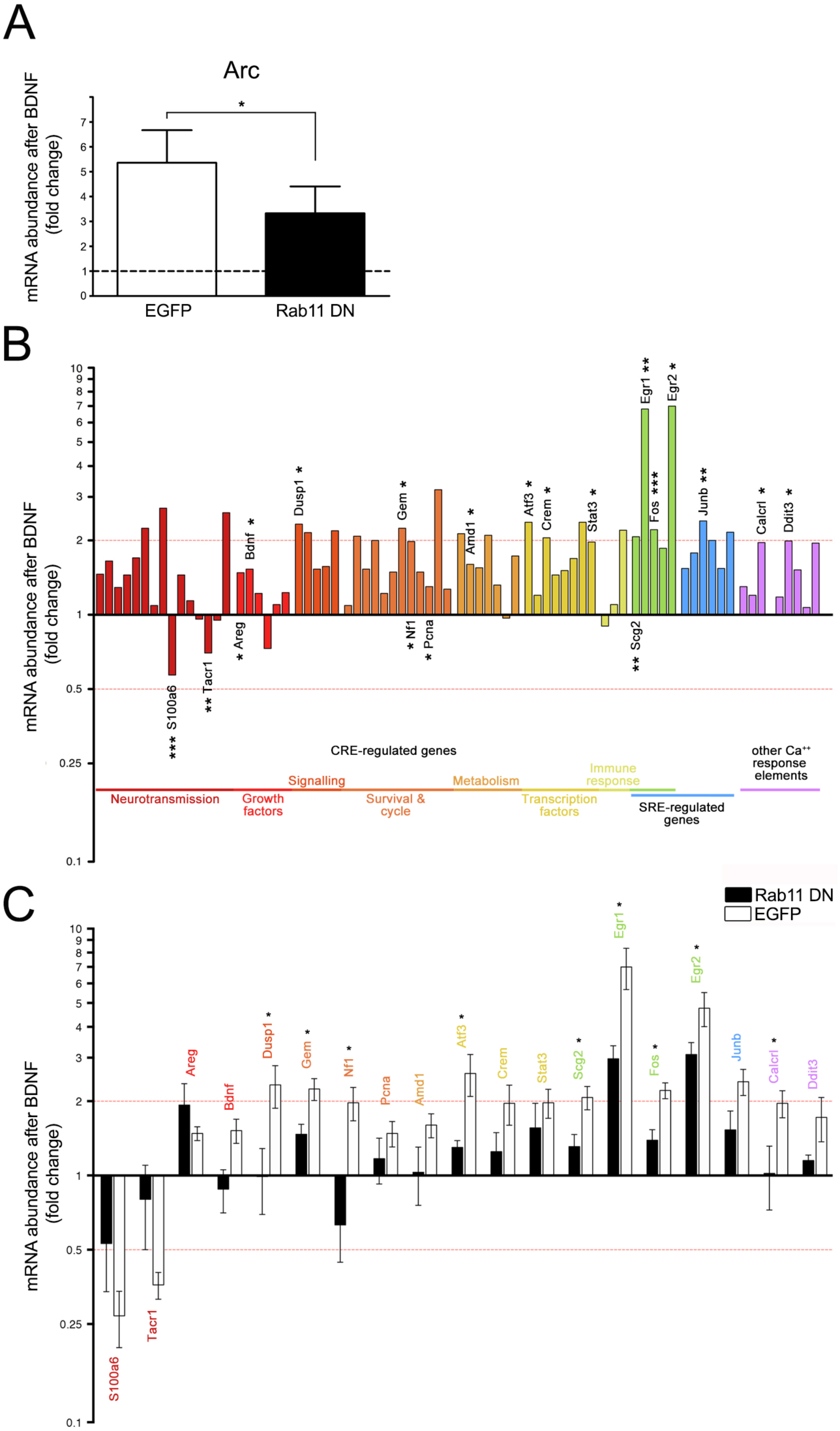
Rab11 activity mediates CREB-regulated gene expression upon stimulation with BDNF. Changes in the expression of the early-response gene *Arc* following 4 hours of BDNF stimulation were studied in neurons expressing control EGFP or dominant negative (DN) mutant of Rab11 and analyzed by using Student’s t test (**A**). Abundance of transcripts was expressed as the fold-change compared to neurons treated with BSA, showing a significant decrease in neurons expressing Rab11DN. To explore BDNF-induced changes in the expression of multiple CREB-regulated genes, we used a PCR array including 84 genes with cAMP, calcium and serum response elements. Reliable data obtained from 76 of the genes in 5 independent experiments were categorized by their annotated functions and plotted as a fold-change in BDNF-stimulated neurons compared to BSA-treated neurons, resulting in significant differences in 19 of these genes (**B**). Although 2 of the studied genes significantly decreased their expression levels upon BDNF stimulation, the remaining 17 were increased. Names and levels of significance are shown for the 19 significantly regulated genes. We performed the same analysis in neurons expressing Rab11DN in parallel to compare the expression levels of these 19 genes. We plotted EGFP and Rab11DN data together for these genes (**C**), finding overall attenuation of the BDNF effect. Moreover, 9 of these genes were significantly different. Names are color-coded following the functional classification. Statistical significance of gene expression data was analyzed by using Student’s t test. Level of significance for the different tests is indicated as follows: *, p < 0.05; **, p < 0.01; ***, p < 0.005; and NS, non-significant.

To further understand the impact of Rab11 in BDNF-induced gene expression, we decided to use a qPCR array for 84 genes regulated by cyclic AMP (CRE), serum (SRE) and Ca^2+^ (CaRE) response elements. We reliably analyzed 76 genes (Supplementary Table S1); 19 of these genes were regulated by BDNF after 4 hours of stimulation (Fig. 5B). The genes significantly modified by BDNF signaling are related to the following neuronal cell responses: neurotransmission (*S100a6, Tacr1*, and *Scg2*); growth factors (*Areg* and *Bdnf*); signaling (*Calcr1* and *Dusp1*); cell cycle, survival and DNA repair (*Gem, Nf1* and *Pcna*); metabolism (*Amd1*); and transcriptional control (*Egr1, Egr2, Atf3, Fos, Crem, Ddit 3, Junb*, and *Stat3*). Then, we analyzed whether the expression of the Rab11DN mutant alters BDNF-induced changes in gene expression. In most cases, hippocampal neurons expressing Rab11DN showed an attenuated response to BDNF, and 9 of these genes were significantly reduced (*Dusp1, Gem, Nf1, Atf3, Scg2, Egr1, Fos, Egr2* and *Calcrl*) (Fig. 5C). Our data indicated that trafficking through the recycling endosomal pathway regulated by Rab11 is crucial for BDNF signaling to define a specific profile of gene expression in hippocampal neurons. Since many of the genes transcriptionally regulated by BDNF signaling and Rab11 activity are related to growth and neuronal plasticity, this finding provides a mechanism explaining how trafficking, signaling and gene expression patterns are integrated towards all-embracing neuronal responses, such as BDNF-induced remodeling of the dendritic tree.

## Discussion

Neurotrophins have multiple roles in the central nervous system, including shaping the morphology of neurons. Through these molecules, neurons integrate local events, such as local remodeling of the cytoskeleton, with long-distance cell responses, including changes in gene expression (Matusica and Coulson, 2014). Whether the endocytic pathway was required for BDNF-mediated nuclear signaling to induce dendritic branching was unclear. We have previously shown that BDNF can regulate the activity and mobility of Rab5 and Rab11 endosomes, and in turn, these GTPases are required for BDNF-mediated dendritic branching (Lazo et al., 2013; Moya-Alvarado et al., 2018). The aim of our study was to determine whether both Rab5 and Rab11 were required for BDNF-induced CREB phosphorylation and transcriptional regulation. We found that both GTPases were required for Erk1/2-dependent CREB nuclear activation and, consistent with the role of the early and recycling pathways in regulating CREB-dependent transcription, Rab11 was also required for BDNF-induced expression changes in genes containing CRE-binding sites. Altogether, our results suggest that endocytic pathways regulate the propagation of BDNF signaling to the nucleus in addition to the local events reported previously (Huang et al., 2013; Lazo et al., 2013; Moya-Alvarado et al., 2018).

An example of the integration between the local and distal effects of neurotrophins is the increase in dendritic complexity triggered by BDNF (Gonzalez et al., 2016). Due to the limited diffusion of this neurotrophin in the extracellular space, BDNF-induced branching is a local process (Horch and Katz, 2002) that involves increased recycling of TrkB receptors, which elevates the local concentration of signaling molecules (Lazo et al., 2013). This mechanism provides signals that regulate the cytoskeleton to induce branching (Sears and Broihier, 2016), allowing elongation of the protruding branch (Ye et al., 2007). However, BDNF-induced dendritic branching depends on the expression of CREB target genes (Finsterwald et al., 2010; Kwon et al., 2011), suggesting a mechanism that integrates local signaling and propagation of the signal towards the nucleus.

Several lines of research suggested that CREB activation requires signaling from intracellular organelles in close proximity with the nucleus. In thyroid primary cells, sustained activation of PKA from the trans-Golgi-network (TGN) is required for CREB activation induced by internalized TSH receptors (Godbole et al., 2017) and sorting of internalized receptor to the TGN-required Rab11 (Nakai et al., 2013; Welz et al., 2014). Furthermore, we have previously shown that a constitutively active mutant of Rab11 allows sensitization to endogenous BDNF, increasing the phosphorylation of CREB (Lazo et al., 2013). This last observation is consistent with BDNF increasing the activity of Rab11 in dendrites as well as the cell bodies of hippocampal neurons and with the requirement of Rab11 activity for increased CREB-dependent transcription (Fig. 5). Moreover, the signaling endosome effector coronin-1 was shown to be required for sympathetic neuronal survival and CREB activation in sympathetic neurons. Coronin-1 allowed the sorting from axonal signaling endosomes to Rab11 recycling endosomes in the neuronal cell bodies, precluding lysosomal sorting and degradation of activated TrkA receptors (Suo et al., 2014). Thus, the Rab11 recycling endosomes constitute a platform for stabilization and integration of signaling receptors, avoiding degradation and allowing intracellular signaling and neuronal function. Signaling endosomes containing activated receptors and signaling molecules help integrate local and distal processes (Zhou and Snider, 2006). These signaling carriers can be diverse in nature, encompassing specialized Rab5-positive early endosomes (Cui et al., 2007; Liu et al., 2007; Wu et al., 2001a), Rab7-positive late endosomes (Deinhardt et al., 2007; Zhou et al., 2012), and Rab11-positive recycling endosomes (Ascano et al., 2009; Barford et al., 2018), among others. Interestingly, the activities of Rab5 and Rab11 are essential for BDNF-induced dendritic branching, and the activity of both GTPases is regulated by BDNF/TrkB signaling (Lazo et al., 2013; Moya-Alvarado et al., 2018).

Wu et al. showed that sustained Erk1/2 activity was critical to induce CREB-dependent gene expression (Wu et al., 2001b), which is in line with BDNF-induced CREB phosphorylation in the nucleus mediated by Erk1/2 in our study. The reduced nuclear pErk1/2 by Rab5 and Rab11DN mutants is therefore consistent with reduced CREB phosphorylation.

Downregulation of CREB nuclear activation was not due to a reduction in the basal levels of the receptor or signaling molecules since 48 hours of expression of the Rab5DN and Rab11DN mutants did not preclude the activation of the receptor or the engagement of the PI3K-Akt and Erk1/2 signaling pathways after 5 minutes of BDNF stimulation. However, two different techniques revealed that sustained activation of Erk1/2 (1-2 hours of BDNF stimulation) was compromised in neurons expressing the mutants. The precise mechanism by which signaling endosomes facilitate the translocation of pErk1/2 to the nucleus needs further investigation; however, based on other cell models (Mebratu and Tesfaigzi, 2009), phosphorylation of Erk1/2 from intracellular organelles may promote its association with a subset of interactors that facilitate its nuclear translocation or prevent Erk1/2 from interacting with proteins that retain it in the cytoplasm. Such a mechanism has been studied for the nuclear targeting of an endosomal E3 ubiquitin ligase in response to increased PKC signaling, reducing the activity of Rab11 with a DNRab11, which abolished nuclear targeting of these endosomal protein (Bocock et al., 2010).

In our study, the gene expression pattern triggered by BDNF signaling was significantly attenuated when Rab11 function was abrogated. The subset of BDNF-regulated genes that depends on endocytic trafficking through the recycling pathway encompasses diverse functional groups involved in metabolism, neurotransmission and regulation of transcription, among others. These genes include early-response genes, such as *Egr1*, *Egr2*, *Fos* and *Arc*, that are important in brain plasticity and memory formation (Heroux et al., 2018; Minatohara et al., 2015).

All these genes, including *Bdnf* itself, are immediate early genes whose transcription is upregulated by activation of CREB in the context of increased neuronal activity (Flavell and Greenberg, 2008; Yap and Greenberg, 2018). Indeed, the overexpression of a constitutively active mutant of CREB led to increased expression of BDNF and BDNF-dependent long-term memory in the hippocampus (Barco et al., 2005). Of note, of the 64 genes analyzed containing CRE binding sites, *Egr1* and *Egr2* were the most highly upregulated by BDNF and also downregulated by reduced Rab11 activity (Fig. 5B and C). The increase in the transcriptional activity of Egr1 was shown to correlate with the activation of different neuronal ensembles involved in learning and memory, anxiety, drug addiction and neuropsychiatric disorders by regulating a variety of genes involved in vesicular release and endocytosis, neurotransmitter metabolism and actin cytoskeleton regulators, among others (Duclot and Kabbaj, 2017). Another TF and immediate-early gene regulated by CREB activation in our study was *c-fos*. Fos forms a heterodimer with Jun to form the activating protein 1 complex (AP1) that regulates activity-dependent transcription in a neuronal subtype-specific manner. Indeed, in addition to binding promoters, AP1 was proposed to bind enhancers, allowing chromatin remodeling and expression of genes in a neuronal population-specific manner (Yap and Greenberg, 2018). Of note, not only activity-dependent genes were upregulated by BDNF. The *Atf3* gene product is a TF involved in neuronal survival, and it was also upregulated by BDNF and downregulated by reducing Rab11 activity (Wu et al., 2013).

One interesting aspect to explore in future studies is the involvement of the BDNF-Rab5/Rab11-Erk1/2-CREB-c-fos-Egr1 pathway in transcription-dependent long-term memory and dendritic morphology maintenance. Specifically, how BDNF and Rab11-mediated CREB activation contributes to a positive feedback loop to regulate activity-dependent transcription should be investigated.

While the role of Egr1-mediated transcription is well documented, the role of Egr2 has been poorly investigated and appears to have opposing functions compared to that of Egr1. Indeed, the behavior analysis of forebrain-specific Cre-mediated Egr2 deletion mice showed improved learning and memory in specific learning and memory tasks (Poirier et al., 2007). In the same line, we also found that BDNF increased the transcription of *Dusp1*and *Nf1*; the products of these genes—a phosphatase and a Ras-GAP, respectively—are known to downregulate Erk1/2 signaling (Lawan et al., 2013; Scheffzek and Shivalingaiah, 2019). However, one of the role associated with Arc in neurons is the endocytosis of AMPA receptors that is required for synaptic depression (Barylko et al., 2018). Thus, our results suggest that the BDNF-Rab5/Rab11-Erk1/2-CREB pathway also triggers genes that downregulate neurotrophic responses, providing a mechanism for homeostatic regulation.

The involvement of BDNF-Rab5/Rab11-Erk1/2-CREB-regulated genes in learning and memory may explain how endosomal system abnormalities and neurotrophin receptor trafficking defects observed in neurodegenerative diseases such as Alzheimer’s disease, Huntington’s disease, amyotrophic lateral sclerosis and Niemann-Pick type C disease are mechanistically linked to dendritic atrophy and synaptic plasticity loss (Brady and Morfini, 2017; Burk and Pasterkamp, 2019; Cabeza et al., 2012; Liot et al., 2013; Xu et al., 2018).

In summary, our results suggest that Erk1/2-dependent nuclear CREB phosphorylation depends on early and recycling endosomal platforms to trigger the genetic program required for dendritic arbor remodeling, among other functions. This work provides new evidence of a mechanistic link between vesicular trafficking and gene expression, reinforcing the possibility of further exploiting the signaling endosomes as a tool to manipulate neuronal physiology.

## Materials and Methods

### Primary culture of hippocampal neurons

Hippocampal neurons were prepared as described previously (Lazo et al., 2013) from E18 rat embryos obtained from the animal facility of the Pontificia Universidad Catolica de Chile and euthanized under deep anesthesia according to the bioethical protocols of our institution. Briefly, hippocampi were dissected and dissociated into single cells in Hank’s balanced salt solution (HBSS, Thermo Scientific); resuspended in Minimum Essential Medium supplemented with 10% horse serum, 20% D-glucose, and 0.5 mM glutamine (MEM/HS); and seeded on poly-L-lysine (1 mg/ml, Sigma) at low density for morphometric and cell biology experiments (7000 cells/cm^2^) or at medium density for molecular biology and biochemical analysis (15,000 cells/cm^2^). After 4 hours, the culture medium was replaced with Neurobasal medium (Invitrogen) supplemented with 2% B27 (Invitrogen) and 0.5 mM glutamine. The proliferation of non-neuronal cells was limited by the use of 0.5 µM cytosine arabinoside (Sigma) beginning at 3 DIV.

### Transfections and infections

To express exogenous proteins, we transfected hippocampal neurons at 7 DIV using Lipofectamine 2000 (Invitrogen) or transduced them with adenoviruses to achieve infection efficiency over 80%. Experiments were performed after 48 hours in complete medium. Plasmids containing pEGFP-C1 are commercially available from Clontech (CA, USA). Rab5 and Rab11 in the pEGFP-C1 vector were gifts from Prof. Victor Faundez (Emory University, USA) and Prof. Rejji Kuruvilla (John Hopkins University, USA), respectively. A non-phosphorylatable mutant of CREB is publicly available from Addgene (#22395). We used replication defective serotype 5 adenoviral vectors to express the EGFP-Rab5 and EGFP-Rab11 DN mutants under the CMV promoter (Ascano et al., 2009; Escudero et al., 2019).

### Western blot

Neurons were washed in PBS and lysed in a solution containing 150 mM NaCl, 50 mM Tris-HCl pH 8.0, 2 mM EDTA, 0.1% SDS, and 1% Triton X-100 supplemented with phosphatase and protease inhibitors and then treated with SDS-PAGE sample buffer. After electrophoretic transfer, nitrocellulose membranes were incubated with the following antibodies: rabbit anti-p490TrkA, 1:1000 (Cell Signaling, cat# 9141); rabbit anti-TrkB, 1:1000 (kindly donated by Dr. Ursula Wyneken, Universidad de Los Andes, Chile); rabbit anti-pAkt, 1:1000 (Cell Signaling, cat# 9271); rabbit anti-Akt, 1:1000 (Santa Cruz Biotechnology, cat# H-136); rabbit anti-pS133 CREB, 1:500 (Cell Signaling cat# 87G3); mouse anti-CREB, 1:500 (Cell Signaling cat# 86B10); rabbit anti-pT202/pY204 ERK1/2, 1:1000 (Cell Signaling cat# 9101); rabbit anti-ERK1/2, 1:1000 (Cell Signaling, cat# 9102); rabbit anti-pY515 TrkB, 1:1000 (Sigma, cat# SAB4300255); rabbit anti-Rab5, 1:1000 (Abcam, cat# ab18211); rabbit anti-Rab11, 1:200 (Zymed/Invitrogen, cat# 71-5300); and mouse anti-βIII Tubulin, 1:1000 (Sigma, cat# T8578). After incubation with the respective secondary antibodies, membranes were developed with SuperSignal Pico West Chemiluminescent Substrate (Thermo Scientific, cat# 34080).

### Immunofluorescence

Neurons were washed in PBS in the presence of phosphatase inhibitors (from here on, the inhibitors were always present in the buffer) and then fixed for 15 minutes in 3% PFA and 4% sucrose dissolved in PBS. Next, the cells were incubated in 0.15 M glycine dissolved in PBS for 10 minutes and then blocked and permeabilized simultaneously by incubation for 1 hour in 4% BSA and 0.5% Triton X-100 in PBS. Incubation with primary antibodies occurred overnight at 4°C. Primary antibodies were diluted in 2% BSA, 0.1% Triton X-100, 0.1 mM CaCl_2_ and 1.5 mM MgCl_2_ dissolved in PBS at the following concentrations: anti-MAP2, 1:200 (EMD Millipore, cat# MAB3418); anti-pCREB, 1:750 (Cell Signaling, cat# 87G3), anti-pERK1/2, 1:400 (Cell Signaling, cat# 9101); and anti-GST, 1:400 (Abcam, cat# ab9085). Then, neurons were washed three times with PBS and incubated for 90 minutes with Alexa Fluor conjugated secondary antibodies 1:400 (Invitrogen) in a solution with a composition identical to that used for primary antibodies. Finally, coverslips were stained for 10 minutes with 1 µg/mL Hoechst 33285 (Invitrogen), washed in PBS, distilled water and mounted with Mowiol 4-88 (Sigma) on microscope slides.

### Analysis of BDNF-induced increases in pCREB and pErk1/2

Hippocampal neurons were starved for 3 hours and stimulated with BDNF for 15 minutes at 37°C. When indicated, neurons were treated from 30 minutes before BDNF stimulation with 30 µM PD98059 (Promega), 10 µM LY294002 (Calbiochem) or 200 nM K252a (Santa Cruz Biotechnologies). At the end of the treatment, neurons were washed in PBS in the presence of phosphatase inhibitors (Thermo Scientific, Cat# 88667), and levels of total and phosphorylated CREB, as well as total and phosphorylated Erk1/2, were evaluated by using phospho-specific antibodies. In experiments involving microscopy, cells were colabeled with Hoechst 33258 (Invitrogen) for 10 minutes. Confocal images were analyzed using ImageJ software, and the integrated intensity (sum of the intensity in a region of interest) was measured within the nuclei. Values are expressed as a BDNF-induced increase in intensity levels with respect to untreated neurons (time 0). When phosphorylation was evaluated by using western blotting, 50 µg of total protein was loaded, and phosphorylation levels were plotted as the intensity of the phosphorylated form with respect to total CREB, TrkB, Akt or Erk1/2 levels.

### Stimulation and measurement of dendritic arborization induced by BDNF

Morphological changes in dendritic arborization induced by BDNF stimulation (50 ng/mL, Alomone) were measured in cultured hippocampal neurons as we previously described (Lazo et al., 2013).

### RNA isolation and quantitative real-time PCR (qPCR)

Total RNA was extracted from hippocampal cell cultures using an RNAeasy Mini Kit from Qiagen and then treated with DNAse I. To analyze BDNF-induced changes in the levels of *Arc*, we synthesized cDNA starting from 1 µg of total RNA, and qPCR was performed with SYBR Green (Applied Biosystems). The custom-made primers used for *Arc* were 5’-GGAGGGAGGTCTTCTACCGT-3’ (fwd) and 5’-CTACAGAGACAGTGTGGCGG-3’ (rev); for *β-actin*, 5’-CCCGCGAGTACAACCTTCT3’ (fwd) and 5’-CGTCATCCATGGCGAACT-3’ (rev); for *Tbp*, 5’-CTGTTTCATGGTGCGTGACGAT-3’ (fwd) and 5’-AAGCCCTGAGCATAAGGTGGAA-3’ (rev); for *Pgk-1*, 5’-TGCTGGGCAAGGATGTTCTGTT-3’ (fwd) and 5’-ACATGAAAGCGGAGGTTCTCCA-3’ (rev). The expression of *β-actin, Tbp* and *Pgk-1* was used to normalize the data between independent experiments. For the array of 84 genes regulated by CRE, SRE and CaRE, we synthesized cDNA starting from 1 µg of total RNA, and we used a Rat cAMP/Ca^2+^ PathwayFinder RT^2^ Profiler PCR Array System following the manufacturer’s instructions (SABioscience, Qiagen, version 4.0, Cat# PARN-066Z). In this case, a combination of five different housekeeping genes (*β-actin, β-2 microglobulin, Hprt-1, Ldha and Rplp1*) was used to normalize the data.

### Statistical analysis

The results are expressed as the average ± standard error of the mean (SEM). Sholl’s analysis curves were compared using two-way repeated-measures ANOVA, followed by Bonferroni’s multiple comparisons post-test to compare all pairs of datasets. Student’s t-test or one-way ANOVA followed by appropriate multiple comparisons test, depending on the number of groups were used when more than two data set required statistical analysis. Details about the specific test used, level of significance and number of replicates are indicated in the respective figure legends. Statistical analyses were performed using GraphPad Prism 5 (Scientific Software).

## Supporting information

Table S1: Genes Read

## Acknowledgements

The authors want to thank Anibal Caceres for help with the qPCR arrays and Professor Giampietro Schiavo (UCL, UK) and Guillermo Moya-Alvarado (Department of Physiology, PUC) for helpful comments on the manuscripts.

## Funding

The authors gratefully acknowledge financial support from FONDECYT grant 1171137 (FB), Basal Center of Excellence in Science and Technology CONICYT (PIA /BASAL PFB 12), Millennium Nucleus (P07/011-F), and the fellowships from CONICYT (AG), Becas-Chile (OML) and The Royal Society (OML).

## Author contributions

AG performed the experiments and drafted the Materials and Methods and figure legends. OML helped with the experimental design, statistics and drafted the article. FB supervised the experimental design and helped draft the article.

## Competing interest

The authors declare no competing financial interests.

