## Supplementary material for "The Rab5-Rab11 endosomal pathway is required for BDNF-induced CREB transcriptional regulation in neurons": Table S1: Genes Read

***Table S1. Genes measured by using the PCR Array***

| Gene | Product | Regulation | Response to BDNF | Function |
|---|---|---|---|---|
| ***Adrb1*** | Adrenoceptor beta 1 | CRE | ns | |
| ***Ahr*** | Aryl hydrocarbon receptor | CRE | ns | |
| ***Amd1*** | Adenosylmethionine decarboxylase 1 | CRE | Increased (*) | Ubiquitous enzyme found in nearly all mammalian cells; may be involved with cell growth |
| ***Areg*** | Amphiregulin | CRE | Increased (*) | May act as a growth factor during central nervous system development |
| ***Atf3*** | Activating transcription factor 3 | CRE | Increased (*) | Expression is associated with neuronal injury |
| ***Bcl2*** | BCL2, apoptosis regulator | CRE | ns | |
| ***Bdnf*** | Brain-derived neurotrophic factor | CRE | Increased (*) | Plays a role in the development of hippocampal long term potentiation; involved in regulation of synaptic plasticity |
| ***Brca1*** | BRCA1, DNA repair associated | CRE | ns | |
| ***Calb1*** | Calbindin 1 | CaRE | ns | |
| ***Calb2*** | Calbindin 2 | CaRE | ns | |
| ***Calcrl*** | Calcitonin receptor like receptor | CaRE | Increased (*) | G-protein coupled receptor for either calcitonin gene-related peptide or adrenomedullin that may play a role in vascular tone |
| ***Calm1*** | Calmodulin 1 | CaRE | ns | |
| ***Calr*** | Calreticulin | CaRE | ns | |
| ***Ccna1*** | Cyclin A1 | CRE | ns | |
| ***Ccnd1*** | Cyclin D1 | CRE | ns | |
| ***Cdk5*** | Cyclin-dependent kinase 5 | CRE | ns | |
| ***Cdkn2b*** | Cyclin-dependent kinase inhibitor 2B | CRE | ns | |
| ***Chga*** | Chromogranin A | CRE | ns | |

| | | | | |
|---|---|---|---|---|
| ***Creb1*** | cAMP responsive element binding protein 1 | CRE | ns | |
| ***Crem*** | cAMP responsive element modulator | CRE | Increased (*) | Interacts with CREB and appears to modulate its response. |
| ***Crh*** | Corticotropin releasing hormone | CRE | ns | |
| ***Ctf1*** | Cardiotrophin 1 | CRE | ns | |
| ***Cyr61*** | Cellular communication network factor 1 | SRE | ns | |
| ***Ddit3*** | DNA-damage inducible transcript 3 | CaRE | Increased (*) | Plays a role in the ER stress response |
| ***Dusp1*** | Dual specificity phosphatase 1 | CRE | Increased (*) | This gene encodes a protein that catalyzes the dephosphorylation and inactivation of MAP kinase, and may be involved in insulin mediated signalling |
| ***Egr1*** | Early growth response 1 | CRE and SRE | Increased (**) | Activates transcription of the LH receptor gene; may be involved in synaptic plasticity during REM sleep |
| ***Egr2*** | Early growth response 2 | CRE and SRE | Increased (**) | Binds DNA; may play a role in learning and long term potentiation |
| ***Eno2*** | Enolase 2 | CRE | ns | |
| ***Fos*** | Fos proto-oncogene, AP-1 transcription factor subunit | CRE and SRE | Increased (**) | An immediate early gene encoding a nuclear protein involved in signal transduction |
| ***Gcg*** | Glucagon | CRE | ns | |
| ***Gem*** | GTP binding protein overexpressed in skeletal muscle | CRE | Increased (*) | GTPase associated to neurons apoptosis and morphology regulation |
| ***Gipr*** | Gastric inhibitory polypeptide receptor | CRE | ns | |
| ***Hk2*** | Hexokinase 2 | CRE | ns | |
| ***Hspa4*** | Heat shock protein family A member 4 | SRE | ns | |
| ***Il6*** | Interleukin 6 | CRE | ns | |
| ***Inhba*** | Inhibin subunit beta A | CRE | ns | |

| | | | | |
|---|---|---|---|---|
| ***Junb*** | JunB proto-oncogene, AP-1 transcription factor subunit | SRE | Increased (**) | Transcription factor; involved in transcriptional regulation |
| ***Jund*** | JunD proto-oncogene, AP-1 transcription factor subunit | CRE | ns | |
| ***Kcna5*** | Potassium voltage-gated channel, shaker-related subfamily, member 5 | CRE | ns | |
| ***Ldha*** | Lactate dehydrogenase A | CRE | ns | |
| ***Maf*** | Avian musculoaponeurotic fibrosarcoma oncogene homolog | CRE | ns | |
| ***Mif*** | Macrophage migration inhibitory factor (glycosylation-inhibiting factor) | CRE | ns | |
| ***Ncam1*** | Neural cell adhesion molecule 1 | CaRE | ns | |
| ***Nf1*** | Neurofibromin 1 | CRE | Increased (*) | GTPase-activating protein that negatively regulates RAS/MAPK pathway. Linked to Neurofibromatosis type 1 |
| ***Nos2*** | Nitric oxide synthase 2, inducible | CRE | ns | |
| ***Npy*** | Neuropeptide Y | CaRE | ns | |
| ***Nr4a2*** | Nuclear receptor subfamily 4, group A, member 2 | CRE and SRE | ns | |
| ***Pck2*** | Phosphoenolpyruvate carboxykinase 2 (mitochondrial) | CRE | ns | |
| ***Pcna*** | Proliferating cell nuclear antigen | CRE | Increased (*) | Processivity factor for DNA polymerase δ. It acts as a scaffold to recruit proteins involved in DNA replication, DNA repair and chromatin remodeling |
| ***Penk*** | Preproenkephalin | CRE | ns | |
| ***Per1*** | Period circadian clock 1 | CRE | ns | |
| ***Plat*** | Plasminogen activator, tissue | CaRE | ns | |
| ***Pln*** | Phospholamban | CRE | ns | |
| ***Pmaip1*** | Phorbol-12-myristate-13-acetate-induced protein 1 | CRE | ns | |
| ***Pou1f1*** | POU domain, class 1, transcription factor 1 | CRE | ns | |
| ***Ppp1r15a*** | Protein phosphatase 1, regulatory subunit 15A | CRE | ns | |
| ***Ppp2ca*** | Protein phosphatase 2 (formerly 2A), catalytic subunit, alpha isoform | CRE | ns | |

| | | | | |
|---|---|---|---|---|
| ***Prkar1a*** | Protein kinase, cAMP dependent regulatory, type I, alpha | CRE | ns | |
| ***Ptgs2*** | Prostaglandin-endoperoxide synthase 2 | CRE | ns | |
| ***Rb1*** | RB transcriptional corepressor 1 | CRE | ns | |
| ***S100a6*** | S100 calcium binding protein A6 (calcyclin) | CRE | Decreased (***) | May function in stimulation of Ca2+-dependent insulin release, stimulation of prolactin secretion, and exocytosis |
| ***Scg2*** | Secretogranin II | CRE and SRE | Increased (**) | Involved in the packaging or sorting of peptide hormones and neuropeptides for secretion. Cleaved to secretoneurin. |
| ***Sgk1*** | Serum/glucocorticoid regulated kinase 1 | CRE | ns | |
| ***Slc18a1*** | Solute carrier family 18 (vesicular monoamine), member 1 | CRE | ns | |
| ***Sod2*** | Superoxide dismutase 2, mitochondrial | CRE | ns | |
| ***Srf*** | Serum response factor | SRE | ns | |
| ***Sst*** | Somatostatin | CRE | ns | |
| ***Sstr2*** | Somatostatin receptor 2 | CRE | ns | |
| ***Stat3*** | Signal transducer and activator of transcription 3 | CRE | Increased (*) | When phosphorylated, it translocate to the nucleus to activate transcription. Plays a roles in cell growth and apoptosis. |
| ***Tacr1*** | Tachykinin receptor 1 | CRE | Decreased (**) | G-protein coupled receptor for substance P. Associated to nitric oxide formation, expresed by cholinergic and nitrergic neurons as well as on smooth muscle cells |
| ***Tgfb3*** | Transforming growth factor, beta 3 | CRE | ns | |
| ***Th*** | Tyrosine hydroxylase | CRE | ns | |
| ***Thbs1*** | Thrombospondin 1 | SRE | ns | |
| ***Tnf*** | Tumor necrosis factor | CRE | ns | |
| ***Vcl*** | Vinculin | SRE | ns | |
| ***Vip*** | Vasoactive intestinal polypeptide | CRE | ns | |

*Level of significance:* *ns = non significant* * = $p < 0.01$ ** = $p < 0.001$ *** = $p < 0.0001$
